# Using a GTR+Γ substitution model for dating sequence divergence when stationarity and time-reversibility assumptions are violated

**DOI:** 10.1101/2020.07.09.195487

**Authors:** Jose Barba-Montoya, Qiqing Tao, Sudhir Kumar

**Affiliations:** Institute for Genomics and Evolutionary Medicine, Temple University, Philadelphia, PA 19122, USA; Department of Biology, Temple University, Philadelphia, PA 19122, USA; Center for Excellence in Genome Medicine and Research, King Abdulaziz University, Jeddah, Saudi Arabia

## Abstract

**Motivation:** As the number and diversity of species and genes grow in contemporary datasets, two common assumptions made in all molecular dating methods, namely the time-reversibility and stationarity of the substitution process, become untenable. No software tools for molecular dating allow researchers to relax these two assumptions in their data analyses. Frequently the same General Time Reversible (GTR) model across lineages along with a gamma (+Γ) distributed rates across sites is used in relaxed clock analyses, which assumes time-reversibility and stationarity of the substitution process. Many reports have quantified the impact of violations of these underlying assumptions on molecular phylogeny, but none have systematically analyzed their impact on divergence time estimates.

**Results:** We quantified the bias on time estimates that resulted from using the GTR+Γ model for the analysis of computer-simulated nucleotide sequence alignments that were evolved with non-stationary (NS) and non-reversible (NR) substitution models. We tested Bayesian and RelTime approaches that do not require a molecular clock for estimating divergence times. Divergence times obtained using a GTR+Γ model differed only slightly (∼3% on average) from the expected times for NR datasets, but the difference was larger for NS datasets (∼10% on average). The use of only a few calibrations reduced these biases considerably (∼5%). Confidence and credibility intervals from GTR+Γ analysis usually contained correct times. Therefore, the bias introduced by the use of the GTR+Γ model to analyze datasets, in which the time-reversibility and stationarity assumptions are violated, is likely not large and can be reduced by applying multiple calibrations.

**Availability:** All datasets are deposited in Figshare: https://doi.org/10.6084/m9.figshare.12594638.

## 1 Introduction

Biological evolution at the molecular level is inherently complex. Nucleotide and amino acid substitution patterns vary from species to species, locus by locus, and over time (Yang, 2014; Arenas, 2015; Nei and Kumar, 2000). Considerable attention has been paid to developing substitution models that better reflect the process of molecular evolution, resulting in increasingly complex, realistic evolutionary models for phylogenomic studies (Yang, 2014; Arenas, 2015). Markov models thoroughly describe the substitution processes that embrace the presence of biased base/amino acid compositions, differences in transition/transversion rates, non-uniformity of evolutionary rates among sites, and differences in substitution patterns among genomic regions (Arenas, 2015; Tao *et al*., 2020).

Widely used substitution models in molecular phylogenetics assume time-reversibility and stationarity of the substitution processes over the whole phylogenetic tree (Yang, 2014; Jayaswal et al., 2011; Galtier and Gouy, 1998). The time-reversibility assumption requires that the instantaneous rate of change from base *i* to base *j* is equal to that of base *j* to *i* (Nei and Kumar, 2000). For large datasets, this assumption is expected to be frequently violated, and an unrestricted model is usually a better fit (Yang, 1994, 2014). While this complexity is well appreciated in molecular evolutionary research, including phylogenetics and systematics, a vast majority of researchers employ a General Time Reversible (GTR) class of substitution models (Fig. 1).

**Fig. 1.**
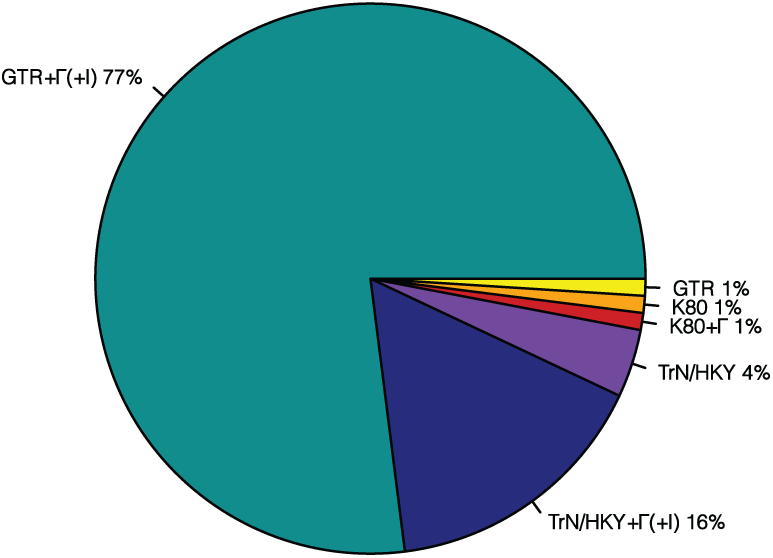
A survey of substitution models selected in 141 research articles that published timetrees in year 2015 – 2017. More than 130 studies (>98%) used models that have more free parameters than the K80 model. All studies assumed stationarity and time-reversiblity of evolutionary processes, with the GTR+Γ and GTR+Γ+I being the most preferred models. K80, HKY, TrN, and GTR represent Kimura-2-parameter (Kimura, 1980), Hasegawa-Kishino-Yano (Hasegawa *et al*., 1985), Tamura-Nei (Tamura and Nei, 1993), and General Time Reversible model (Tavaré, 1986), respectively. Model +Γ(+I) means that either a gamma distribution for incorporating rate variation across sites is used, or a proportion of sites are assumed to be invariant across sequences, or both are used along with the corresponding substitution model.

The GTR model provides for different rates for all the transitions and transversional substitutions as well as the unequal frequency of bases. In addition to time-reversibility, the use of a GTR model in phylogenetic methods, as implemented in most of the software packages, automatically assumes that the substitution process does not change over time; i.e., it is stationary. This translates into assuming that the rates of different types of base substitutions are the same across evolutionary lineages and over time. Violation of this stationarity assumption is evident from differences in base composition across sequences (e.g., Kumar and Gadagkar, 2001; Tamura and Kumar, 2002; Rosenberg and Kumar, 2003; Galtier and Gouy, 1998). Studies of empirical data have shown that unequal base compositions can mislead methods of phylogenetic inference by grouping sequences according to the base compositions instead of their phylogenetic relationships (Lockhart *et al*., 1994; Galtier and Gouy, 1995; Rosenberg and Kumar, 2003; Galen *et al*., 2018). Therefore, many models and methods that avoid the stationarity assumption have been developed (Galtier and Gouy, 1998; Yang, 1994; Blanquart and Lartillot, 2008, 2006; Foster, 2004; Tamura and Kumar, 2002).

Therefore, we expect that the use of the GTR model for substitution rates and phylogenetic inference would cause bias because it is an over-simplification of the correct model. This bias is known to impact phylogenetic inference (Huang *et al*., 2010; Philippe *et al*., 2017; Galen *et al*., 2018; Singh *et al*., 2009), but there is little known about its impact on the estimates of molecular divergence times. In an analysis of bias caused by the use of simple substitution models from the GTR class of models, Tao *et al*. (2020) reported that the complexity of the substitution model has a rather limited biasing effect in empirical data analysis. They found that the actual bias in the time estimates became very small when even a few clock calibrations are applied. The main focus of this study is to examine if this trend holds when we consider datasets that violate the underlying assumptions of time-reversibility and stationarity made in all the current molecular dating analyses.

In the following, we quantified the bias that results from using the GTR substitution model, along with a provision to account for the rate variation among sites by using a Gamma distribution (GTR+Γ), to analyze computer-simulated nucleotide sequence alignments that were evolved without reversibility or stationarity of substitution process. Datasets were simulated for phylogenies in which evolutionary rates varied extensively among lineages with or without autocorrelation among lineages (Tao *et al*., 2019; Rannala and Yang, 2007).

We applied Bayesian and RelTime relaxed clock methods (Tamura *et al*., 2012; Rannala and Yang, 2007) for divergence time estimation. We report that the bias of time estimates caused by the use of a GTR+Γ model with assumptions of stationarity and time-reversibility to analyze datasets that violate these assumptions. Results are presented for analyses using only a root calibration, as well as those in which multiple internal calibrations were assumed to be known.

## 2 Methods

### 2.1 Data simulation

We conducted computer simulations to generate nucleotide sequence alignments in which the substitutional process was reversible (GTR), non-reversible (NR), or non-stationary (NS). All our analyses were conducted by using a model timetree derived from the bony-vertebrate clade in the Timetree of Life (Hedges and Kumar, 2009), from which we sampled 100 taxa (Figs. 2 and 3).

**Fig. 2.**
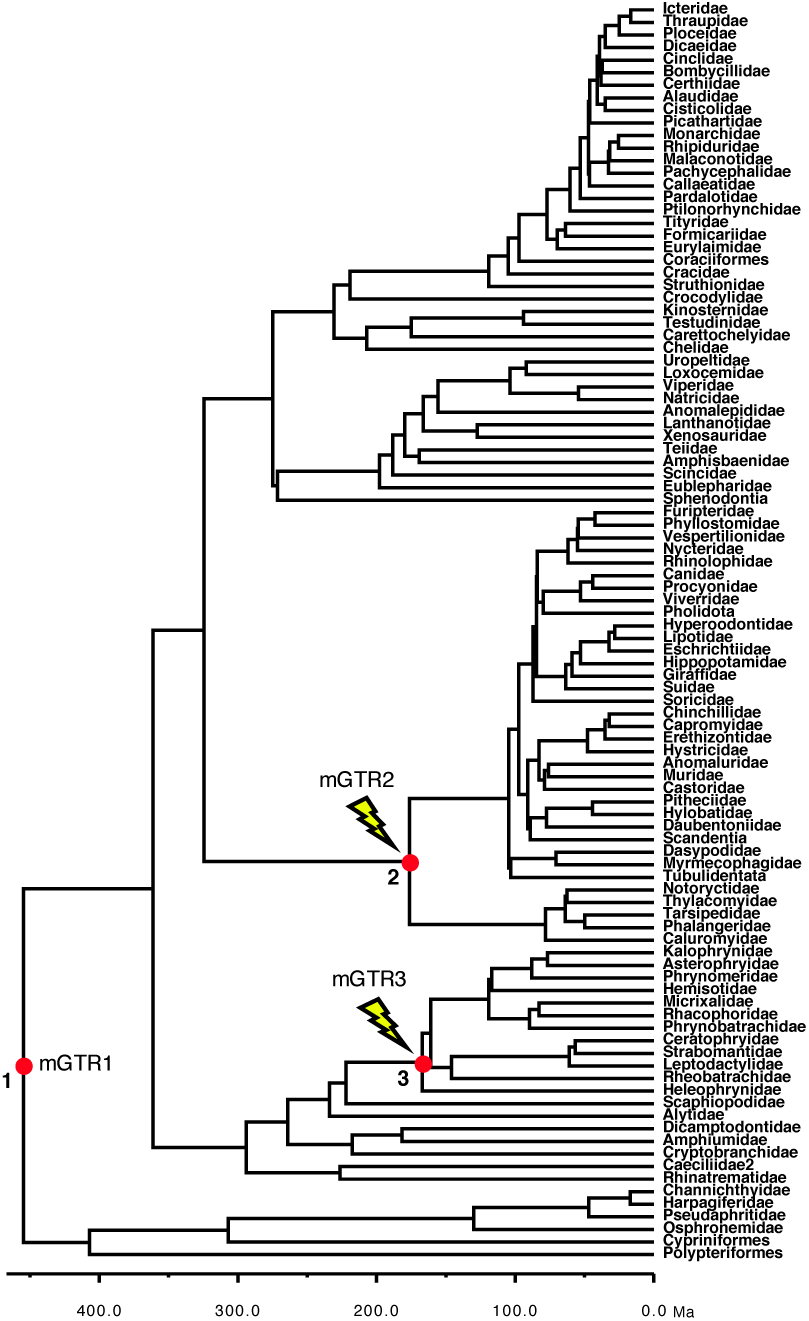
Phylogeny of 100 taxa showing calibrated nodes. The tree has been scaled to time on the basis of time estimates from the Timetree of Life (Hedges and Kumar, 2009). Calibrations are represented for three nodes (red dots). We used a uniform distribution U(min, max) for the three calibrations: (1) root calibration U(444.6, 464.6); (2) Calibration-2 U(166.2, 186.2); (3) Calibration-3 U(157, 177). For the non-stationary alignments, a non-stationary process was added by changing the base composition and rate matrix for two lineages, starting at the ascending branches of node 2 (mGTR2) and node 3 (mGTR3).

**Fig. 3.**
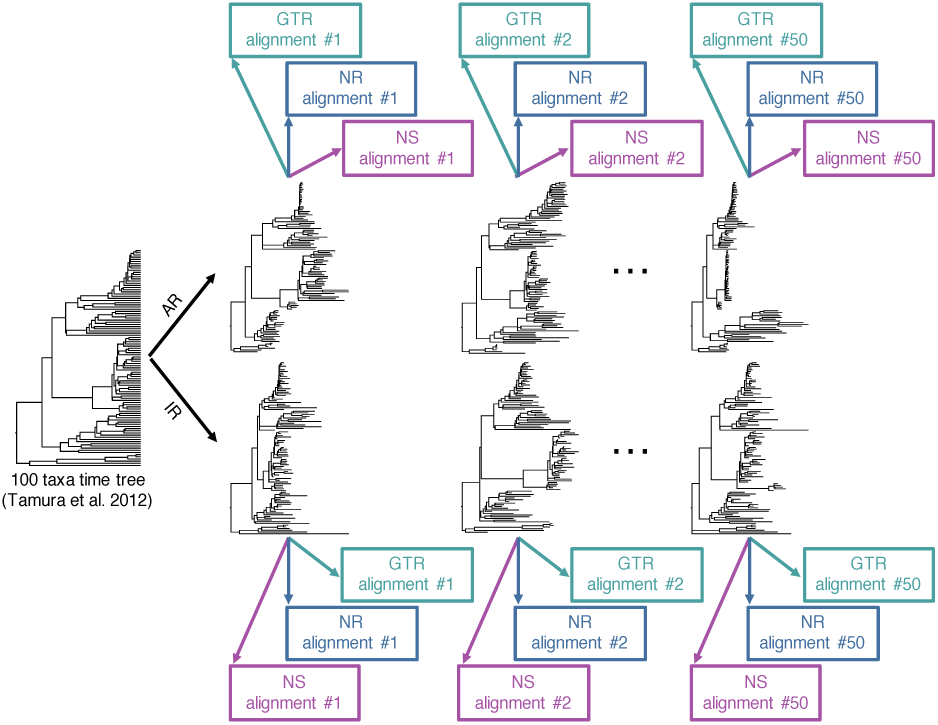
A flowchart showing an overview of the simulation procedure used to generate datasets. We generated 150 alignments of 100 taxa from 50 phylogenies simulated using AR models (50 GTR, 50 NR and 50 NS) and 150 alignments from 50 phylogenies simulated using IR (50 GTR, 50 NR and 50 NS).

We simulated 50 model trees in which the evolutionary rates among branches were autocorrelated (AR datasets) and another 50 in which the rates varied independently (IR datasets); see Tamura *et al*. (2012) for more details. We used INDELible (Fletcher and Yang, 2009) to generate 150 alignments using 50 AR phylogenies such that three datasets were produced from each phylogeny. For one set of data, the nucleotide substitution followed a GTR model with stationarity and reversibility (GTR-AR dataset). In the second, the substitution process was not time-reversible (NR-AR). And, in the third, the substitution process was non-stationary (NS-AR). Similarly, 150 alignments were produced by using IR phylogenies, which resulted in 50 GTR-IR, 50 NR-IR, and 50 NS-IR datasets (Fig. 3).

The GTR alignments were simulated under the GTR+Γ (α = 1.0) model with 1,000 base pairs (bp), a base composition of π_T_ = 0.3, π_C_ = 0.2, π_A_ = 0.3, π_G_ = 0.2, and substitution rate parameters a = 0.2, b = 0.4, c = 0.6, d = 0.8, e = 1.2, f =1. The non-reversible sequences were simulated under the unrestricted model, with 4,000 bp, and substitution rate parameters T→C = 0.1, T→A = 0.2, T→G = 0.3, C→T = 0.4, C→A = 0.5, C→G = 0.6, A→T = 0.1, A→C = 0.2, A→G = 0.3, G→T = 0.4, G→C = 0.5, A→G = 1.

The non-stationary alignments were simulated under the three different GTR+Γ (α = 1.0) models (mGTR1, mGTR2, and mGTR3), with different base composition and rate matrix for different parts of the phylogeny (Fig. 2). Alignments were 4,000 bp long. For mGTR1we used a base composition of π_T_ = 0.3, π_C_ = 0.2, π_A_ = 0.3, π_G_ = 0.2, and substitution rate parameters a = 0.2, b = 0.4, c = 0.6, d = 0.8, e = 1.2, f =1; for mGTR2 we used a base composition of π_T_ = 0.05, π_C_ = 0.45, π_A_ = 0.05, π_G_ = 0.45, and substitution rate parameters a = 0.1, b = 0.2, c = 0.3, d = 0.4, e = 0.6, f =1; for mGTR3 we used a base composition of π_T_ = 0.45, π_C_ = 0.05, π_A_ = 0.45, π_G_ = 0.05, and substitution rate parameters a = 0.15, b = 0.3, c = 0.45, d = 0.5, e = 0.75, f =1.

### 2.2 Estimation of Divergence Times

#### 2.2.1 Bayesian approach

All Bayesian analyses (300 datasets of 100 sequences each) were carried out with the program MCMCTree (Yang, 2007). The correct topology of the 100 taxa tree assumed (Fig. 2) to avoid confounding phylogeny inference errors with divergence time estimation bias. We used the AR model to analyze 150 AR-datasets and the IR model for the 150 IR-datasets; this was done to avoid confounding the bias due to the misspecification of the branch rates model with the bias due to the violation of stationarity and time-reversibility assumptions.

For all the analyses, we assigned to the overall rate (*μ)* a gamma hyperprior G(1, 1) with mean 1/1 = 1 substitutions per site per time unit (100MY) or 10^−8^ substitutions per site per year. To the rate drift parameter (*σ*^2^), we assigned another a gamma hyperprior G(1, 1) with mean 1, allowing large rate variation like the ones simulated here (Tamura *et al*., 2012). The sequence likelihood was calculated under the GTR substitution model with a Γ distribution of site rates (5 categories) (Yang, 1994). The approximate likelihood method (dos Reis and Yang, 2011; Thorne *et al*., 1998) was used in maximum likelihood estimation of branch lengths and the Hessian matrix. The parameters of the birth-death-sampling process were fixed at *λ* = *μ* = 2, and *ρ* = 0.6 (Yang and Rannala, 2006). For each analysis, two runs were performed, each consisting of 5 × 10^6^ iterations after a burn-in of 5 × 10^4^ iterations and sampling every 200, resulting in a total of 5 × 10^4^ samples from the two runs. We checked for convergence by comparing the posterior mean estimates between runs and by plotting the time series traces of the samples. We used two different calibration strategies to investigate the impact of calibrations on time inference: a uniform root calibration only, and a uniform root calibration with another two additional uniform internal calibrations — three uniform calibrations (Fig. 2).

#### 2.2.2 RelTime analysis under relative rate framework

All RelTime analyses were carried out in MEGA-CC software that was prototyped in MEGA X (Kumar *et al*., 2018, 2012). For ensuring a direct comparison, we first used the maximum likelihood (ML) method to estimate branch lengths under the GTR+Γ model with the correct topology by using RAxML (Stamatakis, 2014). Then, each phylogeny with branch length was used to infer a timetree by applying the RelTime method. Timetrees were computed using two different calibration strategies applying uniform constraints: a root calibration only and three calibrations (Fig. 2).

### 2.3 Measurements of accuracies

All comparisons of time estimates (TEs) from simulated data and correct times involved normalized values that were obtained by dividing given node time by the sum of node times in the tree. This procedure avoids normalization biases that may be caused by using any one node as an anchor. The percent difference in time estimates (ΔTE) is the difference between the estimated time and the true time divided by the true time and multiplied by 100. For comparison of model-match and model-mismatch cases, the percent difference in time estimates (ΔTE) is the difference between the estimated and the GTR data TE divided by the GTR data TE and multiplied by 100.

### 2.4 Measurements of coverage probability

We calculated the coverage probability of each node for each dataset. The coverage probability of a node was the proportion of datasets in which the Bayesian highest posterior density intervals (HPDs) or RelTime confidence intervals (CIs) of that node contained the true time. This was done for all datasets with only one root calibration and with three calibrations. True times were normalized to the sum of true times, and lower and upper bounds of HPDs (or CIs) were normalized to the sum of estimated node times.

### 2.5 Measurements of branch length linearity

Because the same phylogeny was used to simulate GTR, NR, and NS datasets, their inferred branch lengths were directly comparable. For each phylogeny simulated under AR or IR rate scenario, we compared the branch lengths inferred using the GTR+Γ model in RAxML for GTR and NR data and GTR and NS data. The coefficient of determination of linear regression through the origin (*R*^2^) was used to determine the linearity of inferred branch lengths. A higher *R*^2^ value indicated a better linear relationship.

## 3 Results and Discussion

We first present results from analyses without applying internal calibrations, which is essential to learn about the intrinsic time structure in the data because calibrations generally impose strong constraints on node ages. We then show results from analyses using a few internal calibrations, which allowed us to examine whether the use of multiple calibrations reduced the bias in times obtained by using the GTR+Γ model.

### 3.1 Impact of violating the time-reversibility assumption

The use of the GTR+Γ model is expected to cause bias in estimating divergence times for the NR datasets because the analysis assumed the time-reversibility of the substitution process (model-mismatch). This bias is explored by comparing the time estimates inferred by using the GTR+Γ model for GTR and for NR datasets simulated using the same phylogeny (50 replicates with autocorrelated and 50 with independent branch rates).

In Bayesian analyses, these estimates showed a high similarity (Fig. 4A and 4B). The mean average percent error (MAPE) was 3.97% when rates were autocorrelated and 1.42% when rates were independent among branches, i.e., the bias is surprisingly small for AR as well as IR datasets. The differences between TEs estimated from GTR and NR datasets (ΔTEs) showed a slightly higher dispersion for AR datasets as compared to the IR datasets. (Fig. 4C).

**Fig. 4.**
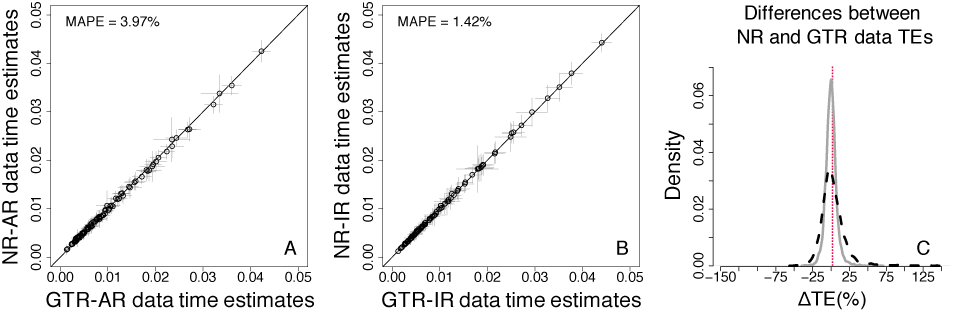
Comparison of Bayesian time estimates obtained by using the GTR+Γ model for analyzing GTR (model-match) and NR (model-mismatch) datasets simulated (**A**) with rate autocorrelation, AR, and (**B**) without rate autocorrelation, IR. Each data point represents the average of normalized times from 50 simulations (±1 SD — grey line). The mean average percent error (MAPE) is shown in the upper left corner of these panels. The black 1:1 line shows the trend if the estimates were equal. (**C**) Distributions of the normalized differences between GTR and NR data TEs for AR (black dashed curve) and IR (grey curve) branch rates. For visual clarity, the distribution in panel C was truncated, removing a few outliers.

Figure 5 compares the dispersions of ΔTEs between the true times and the times obtained when the models matched (panel A) and the times obtained when there was a model mismatch for the NR datasets (panel B). Both of these comparisons show very similar shapes and trends for AR as well as IR datasets. Therefore, the violation of the assumption of the time-reversibility of the Markov process does not seem to have a strong biasing impact in the Bayesian analyses.

**Fig. 5.**
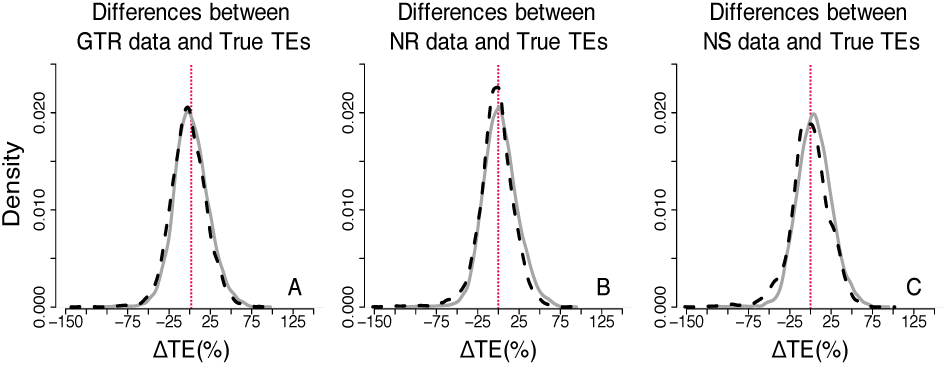
Distributions of the normalized differences between estimated and true node times for GTR, NR, and NS datasets — Bayesian approach (root calibration only). Comparisons of AR (black dashed curve) and IR (grey curve) performance for (**A**) GTR, (**B**) NR and (**C**) NS datasets. For visual clarity, the distribution in panels A, B and C were truncated, removing a few outliers.

Similar results were observed for RelTime estimates. Again, the bias was small (Fig. 6), with MAPE equal to 4.74% when rates were autocorrelated and 1.44% when rates were independent. Moreover, ΔTEs between TEs for NR and GTR datasets showed a low and similar dispersion for both IR and AR data sets (Fig. 6C). However, the RelTime estimates showed greater noise (SDs) as compared to the Bayesian node times. It is not surprising because we assumed correct priors (e.g., tree prior and evolutionary rate model) in Bayesian analyses. In contrast, the RelTime method does not require such prior knowledge in estimating divergence times.

**Fig. 6.**
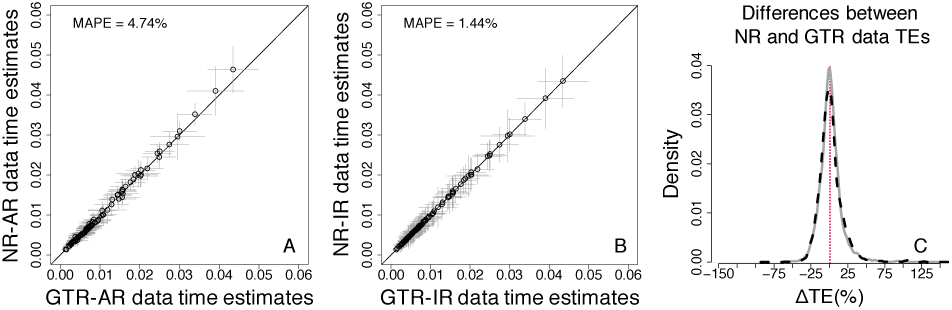
Comparison of RelTime estimates obtained by using the GTR+Γ model for GTR (model-match) and NR (model-mismatch) datasets simulated (**A**) with rate autocorrelation, AR, and (**B**) without rate autocorrelation, IR. Each data point represents the average of normalized times from 50 simulations (±1 SD — grey line). The mean average percent error (MAPE) is shown in the upper left corner of these panels. The black 1:1 line shows the trend if the estimates were equal. (**C**) Distributions of the normalized differences between GTR and NR TEs for AR (black dashed curve) and IR (grey curve) branch rates. For visual clarity, distribution in panel C was truncated, removing a few outliers.

Tao et al. (2020) showed that one reason for the robustness of relaxed clock methods to model misspecification was that the estimates of branch lengths under simple and complex models are often linearly related. In the relative rate framework underlying the RelTime method, divergence times are a function of the ratio of linear combinations of branch lengths. So, we examined the relationship of inferred branch lengths for GTR and NR phylogenies in which a GTR model was used for the inference. We found an excellent linear relationship for an AR and an IR dataset (Fig. 7A and B, *R*^2^ = 0.97 and 0.98 for AR and IR dataset, respectively), which is similar to that reported in Tao et al. (2020). The pattern of linearity of branch lengths was observed across the majority of AR and IR datasets (Fig. 7C). These results indicated similar relative branch lengths were produced when the assumed model matched or did not match the actual evolutionary process, and therefore, similar divergence time estimates. This linear relationship provides a fundamental reason for the results seen for RelTime (Fig. 6) and Bayesian (Figs. 4 and 5) methods.

**Fig. 7.**
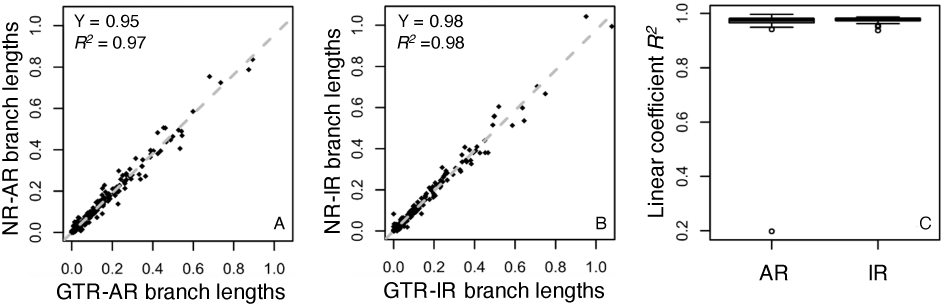
Branch length comparisons for GTR and NR datasets. Branch lengths were inferred by using the GTR+Γ model for (**A**) an AR dataset and (**B**) an IR dataset simulated under the GTR model (x-axis, model-match case) and the NR model (y-axis, model-mismatch case). They all show good linear relationships. The gray dashed line is the best-fit linear regression through the origin. The slope (Y) and coefficient of determination (*R*^*2*^) are shown. (**C**) The dispersion of the linear trends of branch lengths. Boxes show the variation of the coefficient of determination of the linear regression (through the origin, *R*^*2*^) between branch lengths inferred using the GTR+Γ model for 50 GTR and 50 NR datasets simulated under AR and IR scenarios.

### 3.2 Impact of the violation of the non-stationarity assumption

Next, we explored the bias of time estimates caused by the use of the GTR+Γ model to analyze data in which the substitution process was not stationary over time or among lineages (NS datasets). The results of Bayesian analyses showed that the bias on time estimates caused by the use of the GTR+Γ model is again small (Fig. 8 A and B). MAPE between TEs estimated from NS and GTR data was 9.92% when branch rates were autocorrelated and 7.38% when branch rates were independent. The dispersion of node TEs was higher for AR datasets than IR datasets (Fig. 8C). However, overall, the bias is greater for NS datasets than the NR datasets (compare Figs. 4 and 8), with consistent problems observed for some TEs. In particular, TEs from the lineages in which the base composition and rate matrix changed were misestimated. Furthermore, in comparison to NR datasets (Fig. 4C), we found that ΔTEs had a considerably larger dispersion for NS data (Fig. 8C).

**Fig. 8.**
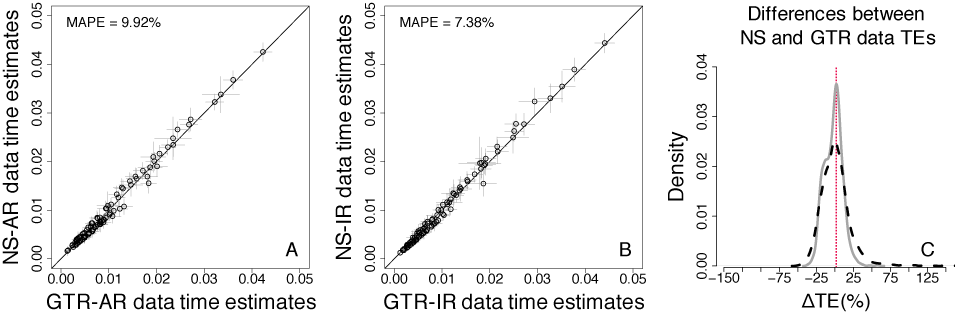
Comparison of Bayesian time estimates obtained by using the GTR+Γ model for GTR (model-match) and NS (model-mismatch) datasets simulated (**A**) with rate autocorrelation, AR, and (**B**) without rate autocorrelation, IR. Each data point represents the average of normalized times from 50 simulations (±1 SD — grey line). The mean average percent error (MAPE) is shown in the upper left corner of these panels. The black 1:1 line shows the trend if the estimates were equal. (**C**) Distributions of the normalized differences between GTR and NS data TEs for AR (black dashed curve) and IR (grey curve) branch rates. For visual clarity, the distribution in panel C was truncated, removing a few outliers.

As for the accuracy of Bayesian times, we found that the distributions of ΔTEs between the estimated and true TEs showed slightly larger dispersion for NS than for NR and GTR datasets under AR and IR models (Fig. 5A vs. C). Therefore, the violation of the assumption of stationary of the substitution process is likely to have a somewhat considerable biasing impact in the Bayesian time inference.

The results of RelTime analyses showed a pattern that resembled the Bayesian methods, as similar time estimates were obtained when the GTR+ Γ model was used on GTR datasets (model-match) and when the GTR+ Γ model was used on NS datasets (model-mismatch, Fig. 9). However, the overall bias on time estimates caused by the use of the GTR+Γ model was slightly larger for RelTime. The MAPE of TEs inferred from NS data was, on average, 12.56% for AR datasets (Fig. 9A) and 11.5% for IR datasets (Fig. 9B). In comparison to NR data (Fig. 5C), the ΔTEs between TEs estimated from NS data and GTR data displayed a larger dispersion (Fig. 9C).

**Fig. 9.**
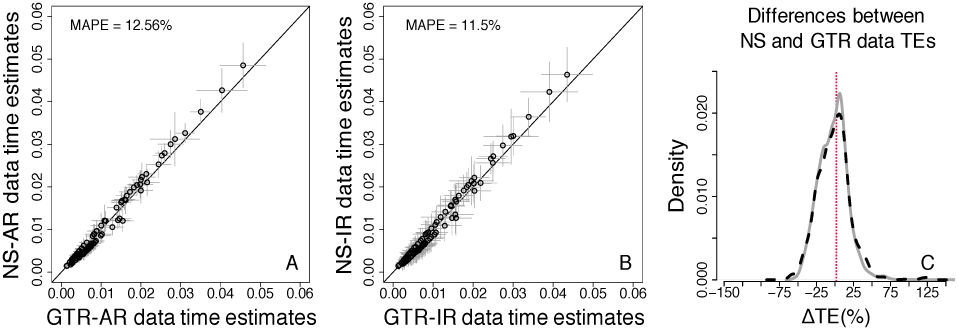
Comparison of time estimates obtained by using the GTR+Γ model for GTR and NS datasets — RelTime approach (root calibration only). (**A**) AR datasets. (**B**) IR datasets. Each data point represents the average of normalized times from 50 simulations (±1 SD — grey line), generated using. The mean average percent error (MAPE) is shown in the upper left portion of each plot. The black line represents equality between estimates. (**C**) Distributions of the normalized differences between GTR and NS data TEs. AR (black dashed curve) and IR (grey curve). For visual clarity, the distribution in panel C was truncated, removing a few outliers.

We also examined the relationship of inferred branch lengths for GTR and NS phylogenies, in which the assumption of stationarity was violated, and a GTR+Γ model was used for branch lengths inference. There was a very high correlation (Fig. 10A and B, *R*^2^ = 0.96 and 0.96 for AR and IR dataset, respectively). However, the linear trend was weaker as compared to that for the NR data and displayed higher variation (compare Figs. 7 and 10). This slightly weaker linear relationship appears to be the reason for the results that higher biased estimates were produced by the use of the GTR+Γ model for NS data than NR data. This is particularly interesting, because figures 8A, 9A, and 10A show very similar trends, indicating that the bias in estimating relative branch lengths is recapitulated in the time estimates produced by both Bayesian and RelTime approaches.

**Fig. 10.**
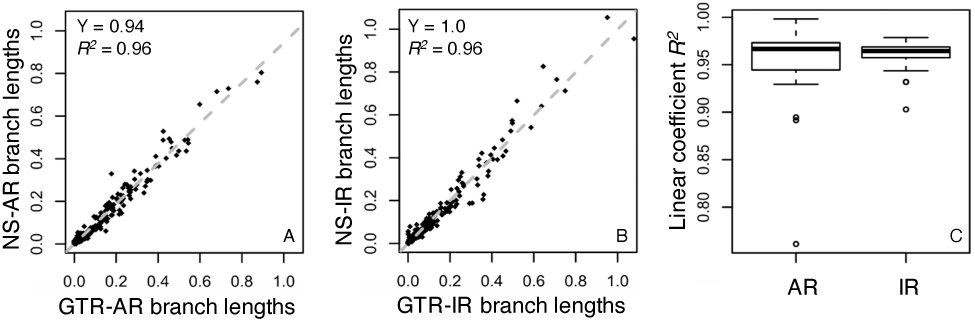
Branch lengths comparisons between GTR and NS data. Branch lengths inferred using the GTR+Γ model for (**A**) an AR dataset and (**B**) an IR dataset simulated under the GTR model (x-axis, model case) and the NS model (y-axis, model case) show a good linear relationship. The gray dashed line is the best-fit linear regression through the origin. The slope (Y) and coefficient of determination (*R*^*2*^) are shown. (**C**) The dispersion of the linear trends of branch lengths. Boxes show the variation of the coefficient of determination of the linear regression (through the origin, *R*^*2*^) between branch lengths inferred using the GTR+Γ model for 50 GTR and 50 NS datasets simulated under AR and IR scenarios.

### 3.3 Improvements offered by the use of calibrations

We also estimated divergence times using two internal calibrations shown in figure 2 to examine whether the use of multiple calibrations constrained the time estimates and reduced the possible error caused by the use of GTR+Γ. As expected, the biased estimates improved when we use calibrations strategically positioned on the nodes that experienced a change in the substitution model and base composition in the phylogeny.

In comparison to analyses with only the root calibration, the MAPE for the NS data was reduced to 8.08% and 6.33% for AR and IR datasets, respectively. For the NR case, the MAPE was reduced to 3.65% for AR datasets, and it remained almost identical (1.46%) for IR datasets (Fig. 11A, B, D, and E). Furthermore, ΔTEs from NR and NS data showed a higher correspondence to those from GTR data (Fig. 11C and F) than in the analyses using a root calibration only (Fig. 4C and Fig. 8C).

**Fig. 11.**
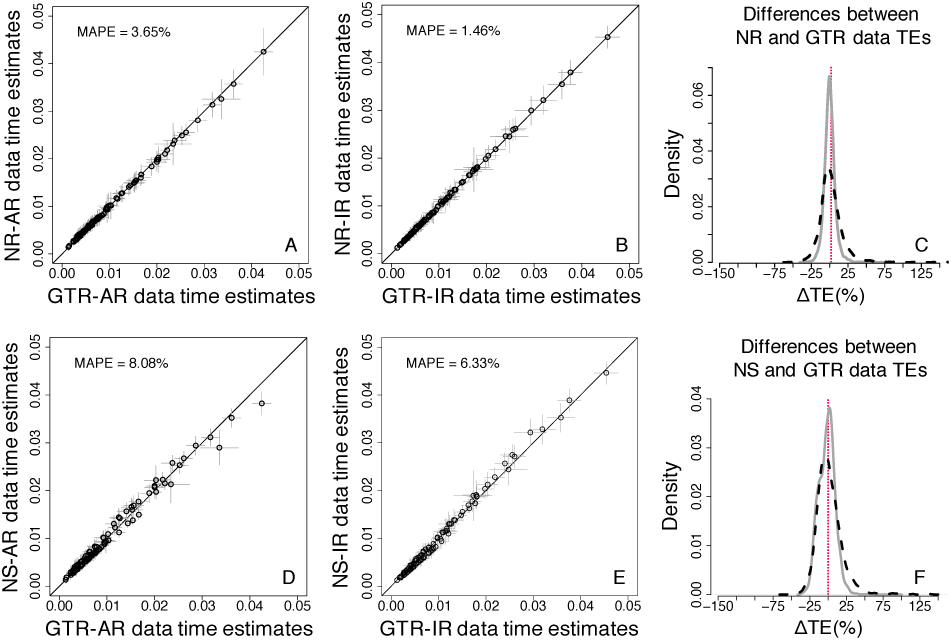
Comparison of time estimates obtained by using the GTR+Γ model for GTR, NR and NS datasets — Bayesian approach (three calibrations). (**A**-**C**) NR datasets. (**D**-**F**) NS datasets. (A, B, D, and E) Each data point represents the average of normalized times from 50 simulations (±1 SD — grey line), generated using. The mean average percent error (MAPE) is shown in the upper left portion of each plot. The black line represents equality between estimates. (**C**) Distributions of the normalized differences between GTR and NR data TEs (F) Distributions of the normalized differences between GTR and NS data TEs. AR (black dashed curve) and IR (grey curve). For visual clarity, the distribution in panels C and F were truncated, removing a few outliers.

The accuracy of Bayesian TEs remained similar when internal calibrations were used (Fig. 12 A-C), although ΔTEs displayed slightly lower dispersion. Overall, ΔTEs showed a high correspondence to those in the analyses using a root calibration only (Fig. 5A-C), excluding NS-AR data, which displayed a significantly reduced dispersion.

**Fig. 12.**
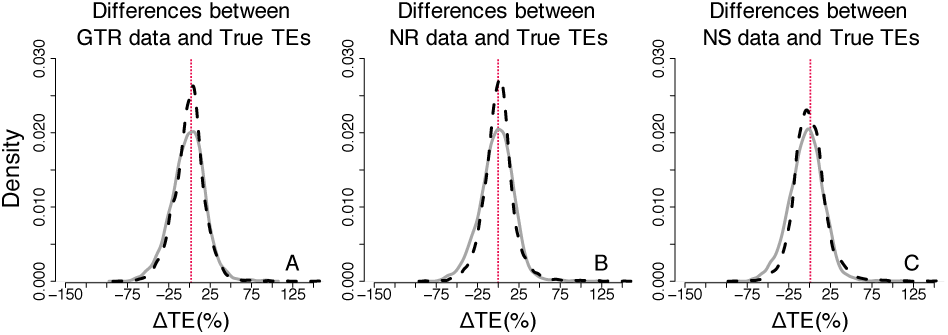
Distributions of the normalized differences between estimated and true times on nodes for GTR, NR, and NS datasets — Bayesian approach (three calibrations). Comparisons of AR (black dashed curve) and IR (grey curve) performance for (**A**) GTR, (**B**) NR and (**C**) NS datasets. For visual clarity, the distribution in panels A, B and C were truncated, removing a few outliers.

### 3.4 Effect of calibrations in the RelTime estimates

Compared to the Bayesian method, we found that the bias on TEs was considerably reduced when RelTime was used, particularly the bias on NR data TEs (Fig. 13A, B, D, and E). When internal calibrations were applied, RelTime TEs inferred from GTR data showed higher similarity to those obtained by using NR and NS data (Fig. 13). The MAPE for NR data was reduced to 1.41% when rates were autocorrelated, and to 1.21% when rates were independent, the MAPE in the NS case was even further reduced to 5.0% and 5.2% for AR and IR data, respectively. Furthermore, ΔTEs from NR and NS data showed a high correspondence to those from GTR data (Fig. 13C and F). These results indicate that the use of interval calibrations can correct bias caused by the use of the GTR+Γ model in analyses of sequence alignments evolved under NS and NR substitution processes. The accuracy of RelTime TEs became higher when internal calibrations were used, as RelTime ΔTEs displayed a reduced dispersion, particularly for NS data (Fig. 14). Although the bias caused by the use of the GTR+Γ model to analyze non-reversible and non-stationary data was reduced, RelTime TEs still displayed a larger dispersion.

**Fig. 13.**
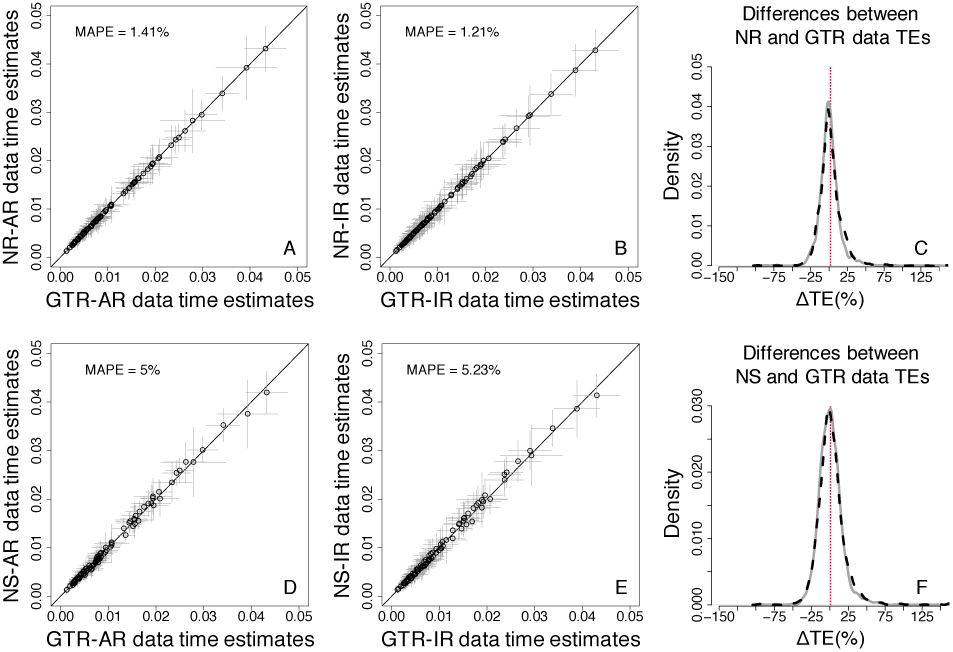
Comparison of time estimates obtained by using the GTR+Γ model for GTR, NR and NS datasets — RelTime approach (three calibrations). (**A**-**C**) NR datasets. (**D**-**F**) NS datasets. (A, B, D, and E) Each data point represents the average of normalized times from 50 simulations (±1 SD — grey line), generated using. The mean average percent error (MAPE) is shown in the upper left portion of each plot. The black line represents equality between estimates. (**C**) Distributions of the normalized differences between GTR and NR data TEs (F) Distributions of the normalized differences between GTR and NS data TEs. AR (black dashed curve) and IR (grey curve). For visual clarity, the distribution in panels C and F were truncated, removing a few outliers.

**Fig. 14.**
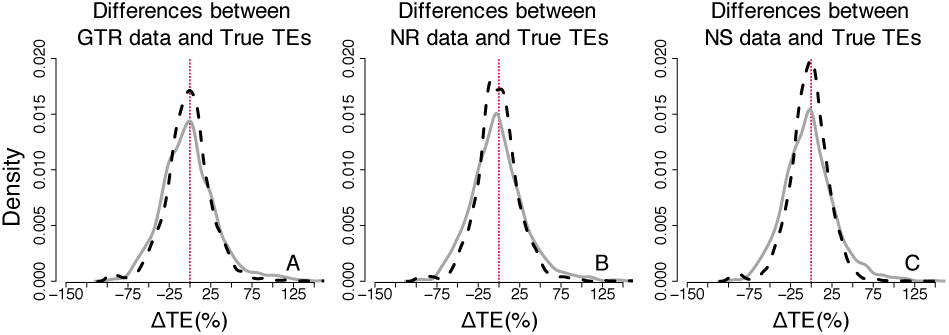
Distributions of the normalized differences between estimated and true times on nodes for GTR, NR, and NS datasets — RelTime approach (three calibrations). Comparisons of AR (black dashed curve) and IR (grey curve) performance for (**A**) GTR, (**B**) NR and (**C**) NS datasets. For visual clarity, the distribution in panels A, B and C were truncated, removing a few outliers.

### 3.5 Evaluation of coverage probability in Bayesian and RelTime approaches

Next, we estimate coverage probabilities that quantified how often the actual node times were contained in the 95% HPDs for Bayesian analyses or 95% CIs for RelTime analyses. We found that Bayesian HPDs obtained using the GTR+Γ model contained the actual node times for the majority of nodes for GTR data (Fig. 15A, median coverage probability = 0.96). This was expected because the used substitution model matched the actual evolutionary process. When the data were simulated under NR and NS models, we also found that HPDs obtained using the GTR+Γ model often included the true times (Fig. 15A, median coverage probability = 0.94 and 0.92 for NR and NS datasets, respectively). More interestingly, distributions of coverage probabilities across all the nodes for NR and NS datasets (model-mismatch cases) were very similar to the one for GTR datasets (model-match case, Fig. 15A).

**Fig. 15.**
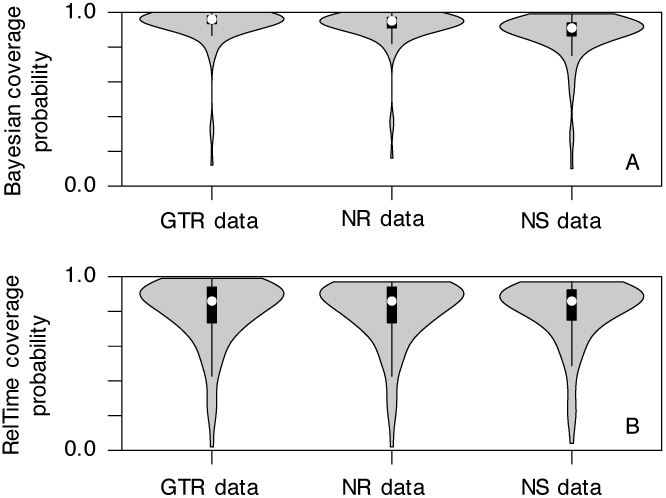
Distributions of coverage probabilities of all the nodes for GTR, NR and NS datasets. Coverage probability of (**A**) Bayesian HPDs and (**B**) RelTime CIs for each scenario is calculated using results of 50 IR and 50 AR simulated datasets obtained using the GTR+Γ model and three calibrations. White dot represents the median value.

Similar distributions of coverage probabilities between matched and mismatched cases were also observed in RelTime analyses (Fig. 15B). These results indicate that the use of the GTR+Γ model on datasets that evolved under much more complex processes is unlikely to impact the estimation of HPDs or CIs, resulting in a consistent conclusion for biological hypothesis testing. However, the overall coverage probability of RelTime CIs was slightly lower than the Bayesian HPDs. It may be because we assumed that correct priors (e.g., tree prior and evolutionary rate model) were known and used them in the Bayesian method, which maximized its performance. In contrast, the RelTime method does not use any such prior knowledge in estimating divergence times.

## 4 Conclusion

In this study, we have analyzed the bias on divergence time estimation caused by the use of the GTR+Γ model to analyze sequence alignments that violate the basic assumptions in phylogenetic analyses: time-reversibility and stationarity of substitution processes. Our results reveal that violating the time-reversibility assumption may only have a limited effect on the accuracy of divergence time estimates. In contrast, the use of sequences with considerable variation in base compositions among sequences, in which the assumption of model stationarity is violated, has a greater effect on the accuracy and precision of divergence time estimates.

Fortunately, we may expect an improvement of accuracy and precision if we use calibrations strategically positioned on the phylogeny. Comparable trends were observed for node time estimates between RelTime and Bayesian analyses, although overall RelTime estimates showed a larger dispersion and higher error. A reason for the better performance of Bayesian analyses is that we used the correct models of rate variation and branching process (birth-death-sampling) for all the analyses. These priors are rarely known beforehand, and RelTime does not require them to be input for the analysis.

Our results are mostly consistent with the conclusion of Tao *et al*. (2020) that show that the complexity of the substitution model has only a modest impact on divergence time estimates. The primary reason for the good performance of the GTR+Γ for analyzing sequences that evolved under non-stationary and non-reversible processes is the high linearity between the branch lengths produced by the mismatched model with those from the correct model. The similar relative branch lengths can be transformed to similar divergence times estimates because the divergence times are a function of the ratio of linear combinations of branch lengths. The present results show that using the GTR+Γ model to analyze sequence alignments, whose basic assumptions are violated, may be sufficient in a majority of time inference tasks. Nevertheless, accounting for time-irreversibility and non-stationarity is still an important aspect of the determination of substitution rates and other phylogenetic inference.

## Acknowledgments

We thank Dr. Marcos Caraballo for technical support.

